# Robustification of RosettaAntibody and Rosetta SnugDock

**DOI:** 10.1101/2020.05.26.116210

**Authors:** Jeliazko R. Jeliazkov, Rahel Frick, Jing Zhou, Jeffrey J. Gray

## Abstract

In recent years, the observed antibody sequence space has grown exponentially due to advances in high-throughput sequencing of immune receptors. The rise in sequences has not been mirrored by a rise in structures, as experimental structure determination techniques have remained low-throughput. Computational modeling, however, has the potential to close the sequence–structure gap. To achieve this goal, computational methods must be robust, fast, easy to use, and accurate. Here we report on the latest advances made in RosettaAntibody and Rosetta SnugDock—methods for antibody structure prediction and antibody–antigen docking. We simplified the user interface, expanded and automated the template database, generalized the kinematics of antibody–antigen docking (which enabled modeling of single-domain antibodies) and incorporated new loop modeling techniques. To evaluate the effects of our updates on modeling accuracy, we developed rigorous tests under a new scientific benchmarking framework within Rosetta. Benchmarking revealed that more structurally similar templates could be identified in the updated database and that SnugDock broadened its applicability without losing accuracy. However, there are further advances to be made, including increasing the accuracy and speed of CDR-H3 loop modeling, before computational approaches can accurately model any antibody.

## Introduction

Antibodies are a crucial component of the adaptive immune system of vertebrates. They are antigen-specific and can be directed towards virtually any antigen to protect us from infections. Their high specificity, in combination with their favorable biophysical properties and pharmacodynamics, have allowed for their development and use as drugs, diagnostics, and research reagents. Antibodies are glycoproteins and are composed of two identical heavy chains and two identical light chains. The isotype is determined by the constant region that dictates effector functions and half life. These constant regions are the same for antibodies of the same isotype. The variable fragments (Fv) on the other hand, are unique to each monoclonal antibody and provide antigen specificity.

Human antibody variable regions consist of a variable light and a variable heavy domain and are extremely diverse, due to V(D)J recombination and somatic hypermutation. These processes result in sequence diversity primarily located in the complementarity determining region (CDR) loops, where the antigen is bound. The CDR 3 loop of the heavy chain (H3) is the most diverse and often particularly important for antigen binding. The remainder of the variable domains is termed framework region and assumes a conserved immunoglobulin (Ig) fold. Antibodies from camelids and cartilaginous fish were found to contain only a variable heavy chain and are referred to as nanobodies, single-domain antibodies, or VHHs.

Antibodies not only play an important role in health and disease, but they are also developed and used as therapeutics. While the availability of sequence information has increased sharply thanks to high throughput sequencing technologies [1], methods for structure determination have remained low throughput. In order to understand the role of antibodies in disease and to efficiently develop drugs, there is a demand for structural information, both for unbound antibodies and for antibodies in complex with their antigens. Computational prediction of these structures is both attractive and feasible because of the relative conservation of the Ig fold across different antibodies [2]. There are several algorithms for antibody structure prediction, such as ABodyBuilder [3], PIGSPro [4], and RosettaAntibody [5]. Across these methods, framework regions are routinely predicted to below 1 Å root-mean-square deviation (RMSD) [6,7], as they pose a simple homology modeling problem wherein a similar structure can be readily identified by a search within a template database. However, the diverse sequences of the CDR loops result in a variety of conformations, making accurate prediction more difficult. All CDR loops, except the H3 loop, fold into clusters of conformations that are termed canonical conformations [8,9]. These loops can be predicted within 1 A RSMD as long as the correct cluster is identified [10,11]. On the other hand, the CDR-H3 loop does not have a limited set of canonical conformations, necessitating *de novo* modeling and resulting in lower accuracy models.

For certain applications, an antibody model suffices, but often there is interest in further downstream modeling, particularly docking against a target antigen. The antigen adds yet another layer of complexity and even more potential for error, especially as the CDR loops can move to accommodate induced-fit binding [12]. Several software packages exist specifically for antibody–antigen docking, including ClusPro [13,14], FRODOCK [15], PatchDock [16], and Rosetta SnugDock [17]. The first three methods are global, rigid-body approaches, adopting different docking algorithms. ClusPro and FRODOCK are fast-Fourier transformation (FFT) based. PatchDock decomposes proteins into geometric patches of hotspots and combines geometric hashing and pose clustering to identify interactions. On rigid targets, for which unbound structures are known, these methods tend to perform well. However, using homology models as input or docking flexible targets remains a challenge. SnugDock was developed to address this challenge. SnugDock is a local, flexible docking method that refines the CDR and V_H_–V_L_ orientation in the context of the antibody–antigen interface. But, to yield low-RMSD models, SnugDock requires an input orientation close to the native, as it is not a global docking approach. A recent assessment revealed that ClusPro was able to find medium quality models (according to the CAPRI definition [18]) less than 40% of time in a global search, while SnugDock found medium quality models 80% of the time on the same set of targets in a local search [19].

Antibody modeling and antibody–antigen docking are fields under active research. Here we report recent developments of RosettaAntibody and SnugDock to improve accuracy of the predicted structures and to make the software more robust and accessible for users and future developers. The template database is now fully automated and can be updated at will, ensuring access to the latest antibody structures in SAbDab [20]. Both RosettaAntibody and SnugDock can now model heavy-chain only antibodies, without any additional flags or specifications. Options for the protocols have been simplified with defaults set based on benchmarks. Constraints have been introduced to improve the quality of models and to allow experimental data to guide modeling. Finally, as these developments were implemented, a set of scientific tests was curated to regularly assess the performance of RosettaAntibody and SnugDock on real-life scenarios.

## Materials and methods

The improvements made to RosettaAntibody and SnugDock are available from Rosetta version 2019.24 on. Note the antibody template database has moved from

`$ROSETTA/tools/antibody` to

`$ROSETTA/main/database/additional_protocol_data/antibody`. The database location can be manually specified with the `-antibody::grafting_database` flag.

### Template database automation

We developed a Python script to query SAbDab, [20] an online antibody database, for the set of sub-3-Å crystal structures. SAbDab pre-processes antibody crystal structures from the PDB and renumbers them according to the Chothia convention [21]. All residue numbers in this manuscript follow the Chothia convention, unless otherwise specified. Based on the information curated by SAbDab, the script then truncates the antibody structures to the relevant structural regions (light chain residues 1–109 and heavy chain residues 1–112). While a crystal structure typically contains a single unique antibody (light chain and heavy chain), there are several structures with multiple distinct antibodies. When multiple chains are present, to avoid ambiguity, we retain the first reported to the SAbDab summary file. If the structure contains only a single light or heavy chain, we retain it. However, if the chain is a single-chain Fv (scFv) (covalently linked light and heavy chain), then it is ignored to limit downstream errors that could arise if the chains are incorrectly assigned. From the truncated structures, sequences are extracted for the regions specified in Table 1 and will be later used in alignments to select structural templates. CDRs containing chainbreaks are omitted during the BLAST database construction. The database is constructed by pooling sequences of the same structural region and length (*e.g.* `database.L1.11` for all sequences of length 11 of the first light-chain CDR) into a single FASTA file, indexed by PDB ID. Each FASTA file is used to build a BLAST database with the `makeblastdb` command. Additionally, the sequences used to construct the database are compiled by structural region and reported to tab-delimited information files for further analysis. Finally, average B-factors for all atoms in each CDR loop and V_H_–V_L_ relative orientation metrics are computed, so these values can later be available to quality filters.

**Table 1.** Structural region to sequence mapping for RosettaAntibody.

| Region<sup>a</sup> | Definition<sup>b</sup> |
| --- | --- |
| CDR L1 | 24–34 |
| CDR L2 | 50–56 |
| CDR L3 | 89–97 |
| FRL<sup>c</sup> | 10–23 |
|  | 35–39 |
|  | 46–49 |
|  | 57–66 |
|  | 71–88 |
|  | 98–104 |
| CDR H1 | 26–35 |
| CDR H2 | 50–65 |
| CDR H3 | 95–102 |
| FRH<sup>c</sup> | 10–25 |
|  | 36–39 |
|  | 46–49 |
|  | 66–94 |
|  | 103–109 |
| Orientation | L5–L104<br>H5–H109 |

<sup>a</sup>CDR and framework region definitions in RosettaAntibody. These definitions are used to extract sequences and templates for the homology modeling database. In a modeling task, templates are selected for each region and combined to produce the initial homology model. <sup>b</sup>The definitions are based on the Chothia numbering convention, but are modified for use in RosettaAntibody. <sup>c</sup>The FRH and FRL definitions do not exactly complement the CDR definitions as there are additional (non-CDR) loops that are excluded from the frameworks.

The automated database can be generated by running the `create_antibody_db.py` script. A comparison of the last version of the manual template database and the first version of the automatic template database is presented in the results section.

### Enabling nanobody—antigen docking

In the grafting step of RosettaAntibody we removed the requirement for a light chain. Using the flag `—vhh_only` it is now possible to produce heavy-chain only antibody models. Within SnugDock, we now apply a hierarchical kinematic representation (referred to as a `FoldTree`) of the antibody–antigen complex by taking advantage of “virtual” residues. In Rosetta, such residues are ignored during energy calculations, but can be used to describe translations and rotations. Throughout SnugDock, a single, “universal” `FoldTree` permitting both V_H_–V_L_ and antibody-antigen docking motions is implemented as described in Figure 1.

**Fig 1.**
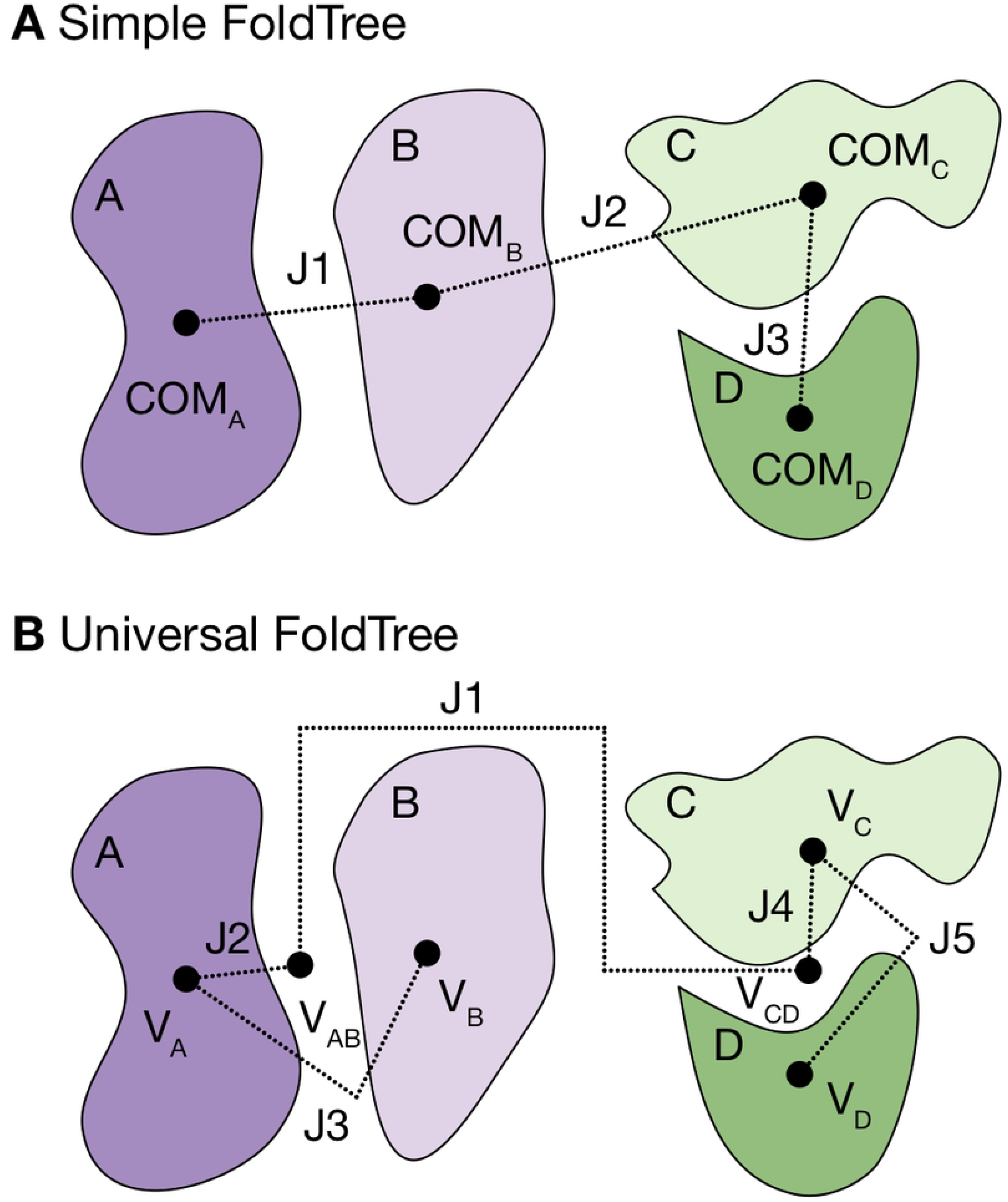
Comparison of a default FoldTree versus one permitting multi-body docking. Proteins are shown as blobs and labeled A, B, C, and D, jumps (describing relative translations and rotations) are labeled as “J” followed by the number, indicating the order of appearance in the `FoldTree`, and the jumps connect either protein centers of mass (COMs) or virtual residues (V). **A:** With the default, linear `FoldTree`, protein A can dock to the BCD protein complex, the AB complex can dock to the CD complex, and the ABC complex can dock to protein D, but protein B cannot move independently. **B:** In the new hierarchical FoldTree, subcomplexes of interest (*e.g.* antibody chains or antigen chains) are grouped by virtual residues, such that the resultant `FoldTree` permits relative motions. In the exemplary implementation, virtual residues are positioned at the individual chain COMs and complex COMs. Docking across individual chain COMs allows for motions within complexes, whereas docking across complex COMs allows for cross-complex docking.

### Simplified options, new filters, and new constraints

Improvements were made to the options, filters, and constraints within RosettaAntibody and SnugDock. Briefly, we reduced the number of options required to be set by the user in both protocols by setting optimal defaults based on our benchmarking simulations. For the homology modeling stage of RosettaAntibody, we implemented new filters as command-line options to permit the exclusion of specific template PDB files or of cases where the template and query have sequence mismatches involving proline residues in the CDR loops (see Results). Finally, we implemented an automatic glutamine-glutamine (Q–Q) hydrogen bonding constraint in the CDR-H3 loop modeling stage of RosettaAntibody and in SnugDock.

The Q–Q constraint is described by a flat harmonic potential:

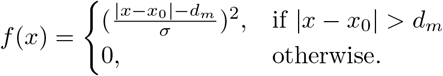

Here, *x* is the distance between the donor and acceptor heavy atoms, *x*_0_ is the mean observed distance in our antibody database, *d_m_* is the minimal difference at which the penalty will be applied, and *σ* is the observed standard deviation. There are two possible hydrogen bonds between Gln 39 of the heavy chain and Gln 38 of the light chain. We measured the donor–acceptor distances for all antibodies in our updated antibody database that contain the relevant Gln residues. The fit is shown in Figure S1. The distances between the N and O atoms yielded *x*_0_ = 2.91 A and σ = 0.23 A. The *d_m_* value is chosen to be 0.5 *σ*, such that there is no penalty in being within half a standard deviation of the mean and there is a penalty of 0.5 REU at one standard deviation.

### New loop modeling and scientific benchmarks

The final set of improvements to RosettaAntibody and SnugDock brought a new, fragment-based loop modeling approach and new scientific benchmarks to both methods. Both the new loop modeling approach (Pan, X. *et al.*) and the scientific benchmarking framework (Leman, J. K. *et al.*) will be fully detailed in other publications that are currently in preparation.

## Results

### Scientific benchmarking

In the process of developing RosettaAntibody and SnugDock, a series of scientific benchmarks were developed. Scientific benchmarking complements other forms of software testing, such as integration and unit tests, by assessing Rosetta’s performance on a diverse set of relevant modeling challenges. A single scientific benchmark consists of a full simulation, whereas unit tests focus on individual functions and integration tests assess exact changes in output. A scientific benchmark is considered successful if the performance is within a certain threshold, usually set by a prior publication. We created three scientific tests for RosettaAntibody and SnugDock. The tests are based on previously published datasets and run regularly on a webserver (https://benchmark.graylab.jhu.edu). There are two RosettaAntibody tests: grafting and loop modeling. While grafting is a fast process (≤ 10 mins per model), CDR-H3 loop modeling is time consuming, so the tests were split based on their runtime. The grafting test runs the `antibody` executable for 48 targets (listed in the Appendix S1), originally described in [22], and it evaluates the RMSDs between the grafted models and the native crystal structures over all antibody structural regions (Table 1). The CDR-H3 loop modeling test runs the `antibody_H3` executable for a six-target subset of the Marze *et al.* set [22], ranging from easy to difficult, and it evaluates the RMSDs between the models and crystals for the CDR-H3 loop. There is a single SnugDock test that is run on six targets (again ranging from easy to hard) and assesses the interface RSMD between the modeled complexes and the corresponding crystal structures.

### Template database improvements

A homology modeling method, such as the grafting stage of RosettaAntibody, is highly dependent on the structural database it samples for templates. A database with inadequate template coverage will result in poorer modeling outcomes. In the most recent CAPRI assessment [23], we were tasked with modeling two camelid antibodies but could not find suitable non-H3 CDR loop templates in the RosettaAntibody database. Further investigation revealed that the database was outdated and contained artifacts due to its manual curation, a consequence of its initial development in 2008. At that point in time, antibody structures were few and antibody structure databases with consistent numbering schemes, such as IMGT [24], SAbDab [20], and abYsis [25], were not yet developed or, in the case of IMGT, did not use a numbering scheme compatible with RosettaAntibody.

Now we have a new antibody template database that can be automatically generated and updated with the `create_antibody_db.py` script. Table 2 shows the increase in available templates following the update and Figure 2 shows the increase in unique sequence templates for each structural region, which is in the range of 15–49%. For the overlapping portion of the two databases (identical PDBs), we compared the template structures and sequences to ensure that no drastic changes had occurred. We found approximately 1–2% of the overlapping templates mismatched at the sequence level, depending on the structural region.

**Fig 2.**
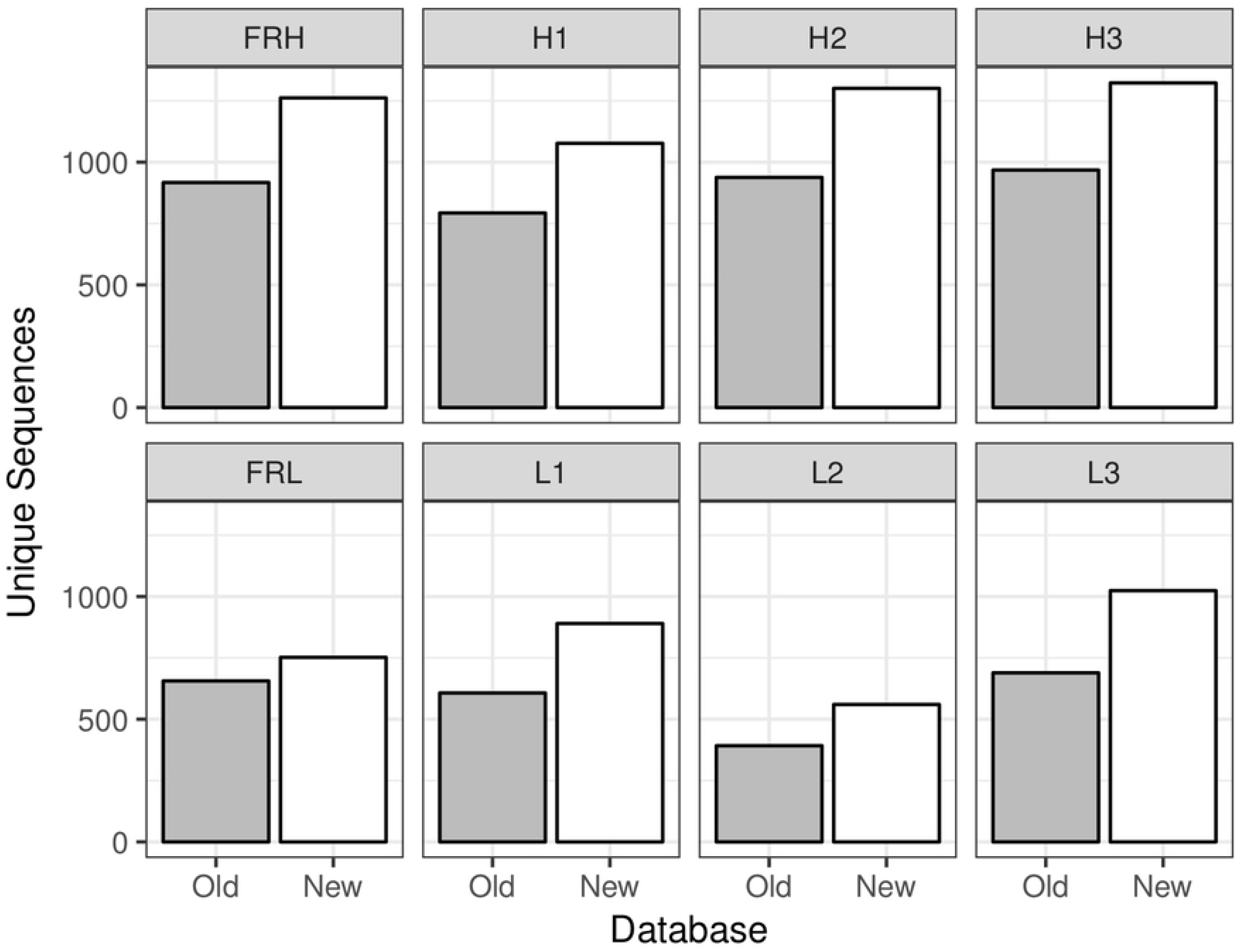
New database increases unique sequence count for all regions. Comparison of counts of unique sequences for each structural region in the old manually curated database (gray, last updated May, 2017) and new automatically generated database (white, last updated June, 2019).

**Table 2.** More templates are available for all structural regions in the new database.

|  | Old Count | New Count | Overlap |
| --- | --- | --- | --- |
| All CDRs | 1,902 | 2,611 | 1,560 |
| FRH | 1,785 | 2,390 | 1,427 |
| FRL | 1,577 | 2,832 | 1,111 |
| Orientation | 1,003 | 1,721 | 749 |

Comparison of the template count between the last iteration of the manual database and the first iteration of the automatic database (February 15th, 2019). Template counts for each region are shown as well as the “overlap” or number of shared templates between the two databases. Additionally, some sequences in the old database do not appear in the new database because it has more stringent quality criteria.

Investigating the individual cases that differed revealed three general trends. One set of cases arose when multiple antibodies were present in the same PDB asymmetric unit and different antibodies were selected from the multiple possibilities. In another set of cases, PDBs were omitted in the new database because they did not adhere to the new quality criteria. *I.e.* these structures had missing atoms or non-realistic C–N distances in critical regions. Finally, the most prevalent set of cases revealed differences in the heavy-chain framework or CDR-H2 loop because the numbering schemes differed as the old database had incorrectly numbered a few highly variable loops, possibly because the regular expressions failed to account for edge cases such as engineered antibodies. In the new database, numbering errors are avoided because structures and sequences are derived from Chothia-numbered PDB files that have been numbered more accurately. [26]

### RosettaAntibody improvements

Improvements to RosettaAntibody affected either the grafting or CDR-H3 loop modeling stage.

#### Grafting with an expanded database and filters

In the grafting stage, RosettaAntibody benefited from the new template database and new filters. Figure 3 shows a direct comparison of the grafted models for 48 target antibody sequences, previously described in [22]. We omit 3MLR due to its atypical CDR L3 loop. We report RMSDs of the loops and framework regions, as well as Orientational Coordinate Distance (OCD), a measure of the relative orientation between the heavy and light chain [22]. In general, we found that the new database produces lower-RMSD grafted models for 53.5–55.0% of target regions. This set of grafting targets was implemented as an automatic scientific benchmark.

**Fig 3.**
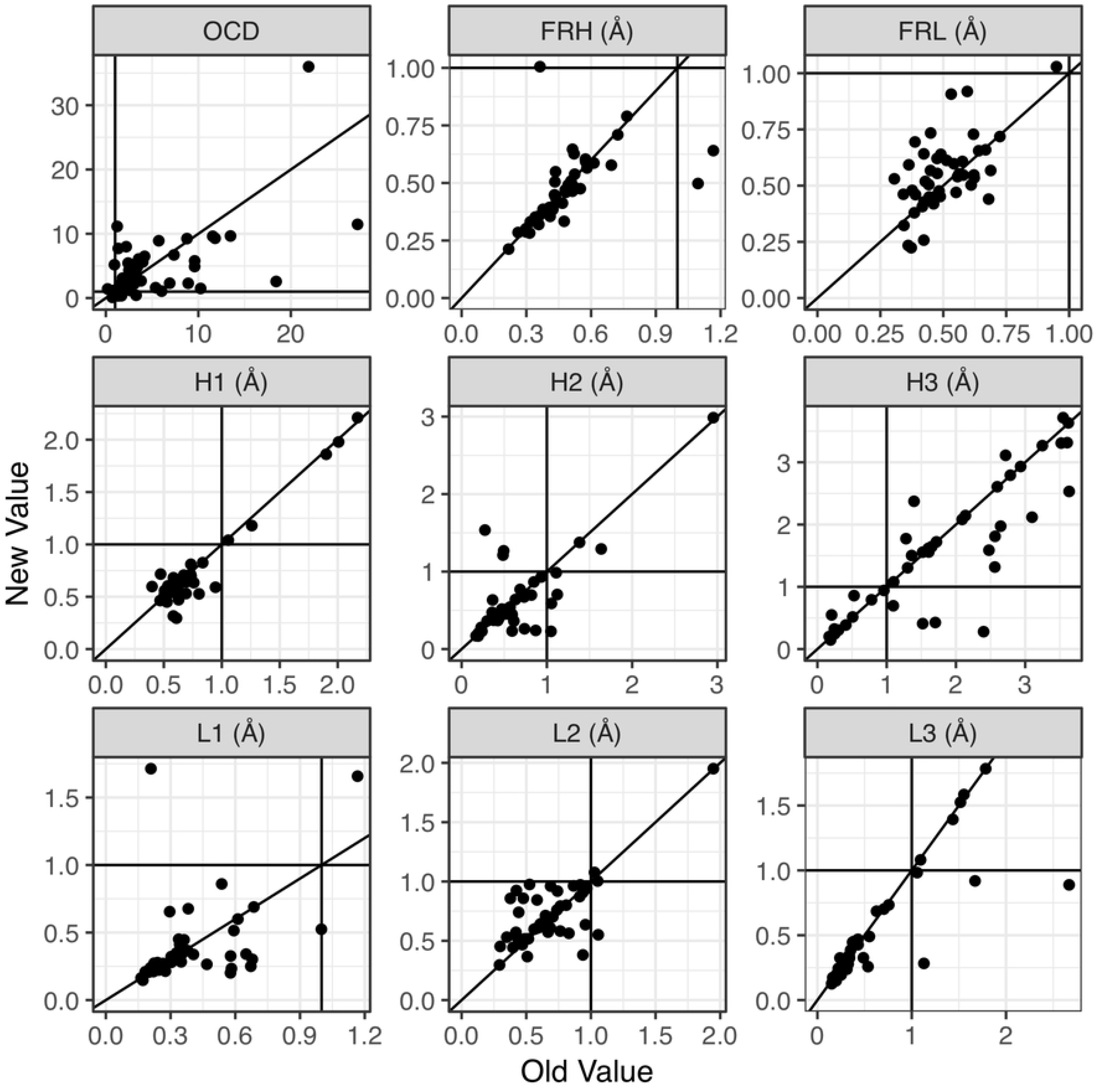
Comparison of grafted model metrics produced by the new and old databases. Plots show values for structural metrics (either OCD or RMSD), comparing the grafted models to the native structures as generated using either the old database (manually curated, x-axis) or new database (automatically curated, y-axis). Solid lines indicate either 1 A or 2 OCD and the diagonal (*i.e.* expected performance if there is no change). The OCD is a measure of the heavy-light chain orientation, previously described in [22]. The new database slightly improves the performance of the grafting step of RosettaAntibody, with 55% of CDR loops and 53.5% of FRs having lower RMSDs.

Following template selection (based on sequence similarity) potential templates are then filtered based on certain criteria. We introduced a PDB ID and a proline filter to improve the selection process for non-H3 CDR templates. The PDB ID filter excludes a particular PDB from the template set, *e.g.* `-antibody:exclude_pdb 1AHW`. This is useful for benchmarking; if the query sequence has a known structure, then it can be excluded from the template set. The proline filter ensures that prolines match between the query sequences and template structures. Prolines occupy a distinct region of Ramachandran space, but the current template selection approach, BLAST, uses either the BLOSUM62 (for framework alignments) or PAM30 (for CDR alignments) matrix and does not sufficiently penalize proline mismatches. While the filter eliminated proline–non-proline mismatches between template and query sequences, it did not demonstrate a concrete improvement in terms of loop RSMD (Figure S2).

#### CDR-H3 loop modeling with fragments and V_H_–V_L_ refinement with constraints

In the CDR-H3 loop modeling stage, we simplified options and introduced a new fragment-based loop modeling method. The options system permits users to pass values to compiled Rosetta binaries via flags on the command line. To configure the CDR-H3 loop modeling stage of RosettaAntibody, a user previously had to specify the loop modeling method, its settings, and custom constraints to maintain the Q–Q bond at the V_H_–V_L_ interface, if present. This constraint is now automated and included by default. The legacy options `-cter_insert`, `-flank_residue_min` (bool), `-bad_nter (bool)`, `-idealize_h3_stems_before_modeling (bool)`, `-remodel (string)`, and `-refine (string)` have been completely removed. C-terminal H3 insertions can now be accomplished via fragment-based kinematic loop closure (KIC). We no longer minimize flanking residues during loop modeling or manually adjust CDR-H3 loop dihedral angles, bond angles, and bond lengths, as this does not affect performance. Finally, the `remodel` and `refine` options are removed. These options previously set the loop modeling algorithm, but the loop modeler is now fixed to be KIC, as it has been shown to be the most accurate approach within Rosetta [27]. Furthermore, by refactoring the code to use the newly developed `LoopModel` class, all other loop-related options are by default set to reasonable values, so it is no longer necessary for the user to configure loop-modeling options, although the possibility remains. In sum these efforts have reduced the number of options required to configure RosettaAntibody from approximately 30 to 5 Appendix S2.

We implemented a new fragment-based loop modeling approach as it was found that fixing sub-regions of loops to match the structures of short fragments (either of length three or nine residues) of similar sequence improved both the fraction of sub-ÅA models and the RMSDs of near-native models (Pan, X., personal communication). Fragments were selected via the fragment picker on the Robetta server [28]. The new loop modeling method was tested on 49 antibody targets from Marze *et al.* [22] and showed no difference in performance when compared to the standard approach. In particular, we expected the use of structural fragments to enhance sampling during loop modeling and lower the minimum RMSD observed across all models. Instead we observed a slight worsening of this metric in the fragment-based models (Figure 4A). As this lack of improvement may have been caused by the highly unique nature of the CDR-H3 loop, we sought to quantify the structural similarity between both protein and CDR-H3 loops and the fragments used in modeling. We investigated the structural similarity between the fragment sets picked for loop modeling and the corresponding target antibody CDR-H3 loop or other (non-antibody) protein loop. For each loop and each possible window of size three or nine residues, the fragment picker selected two hundred selected fragments. These fragments and their corresponding loop segments were compared by measuring the average difference in the backbone dihedral angles as a chord distance (originally defined by Dunbrack and North [8]). We found that non-antibody protein loops were more likely to have near-native fragments identified by the picker than antibody CDR-H3 loops (Figure 4B). This was due to one of two possibilities: (1) either structurally similar fragments exist and the fragment picker cannot identify them for antibody CDR-H3 loops or (2) the fragments do not exist. Considering that the fragment picker tends to perform well across a diverse set of targets [28] and previous observations that antibody CDR-H3 loops have fewer structurally similar fragments in the PDB than than other protein loops [29], we concluded that the latter is most likely and the lack of structural similarity between fragments and CDR-H3 loops can explain the inability of fragment-based loop modeling to improve CDR-H3 loop models.

**Fig 4.**
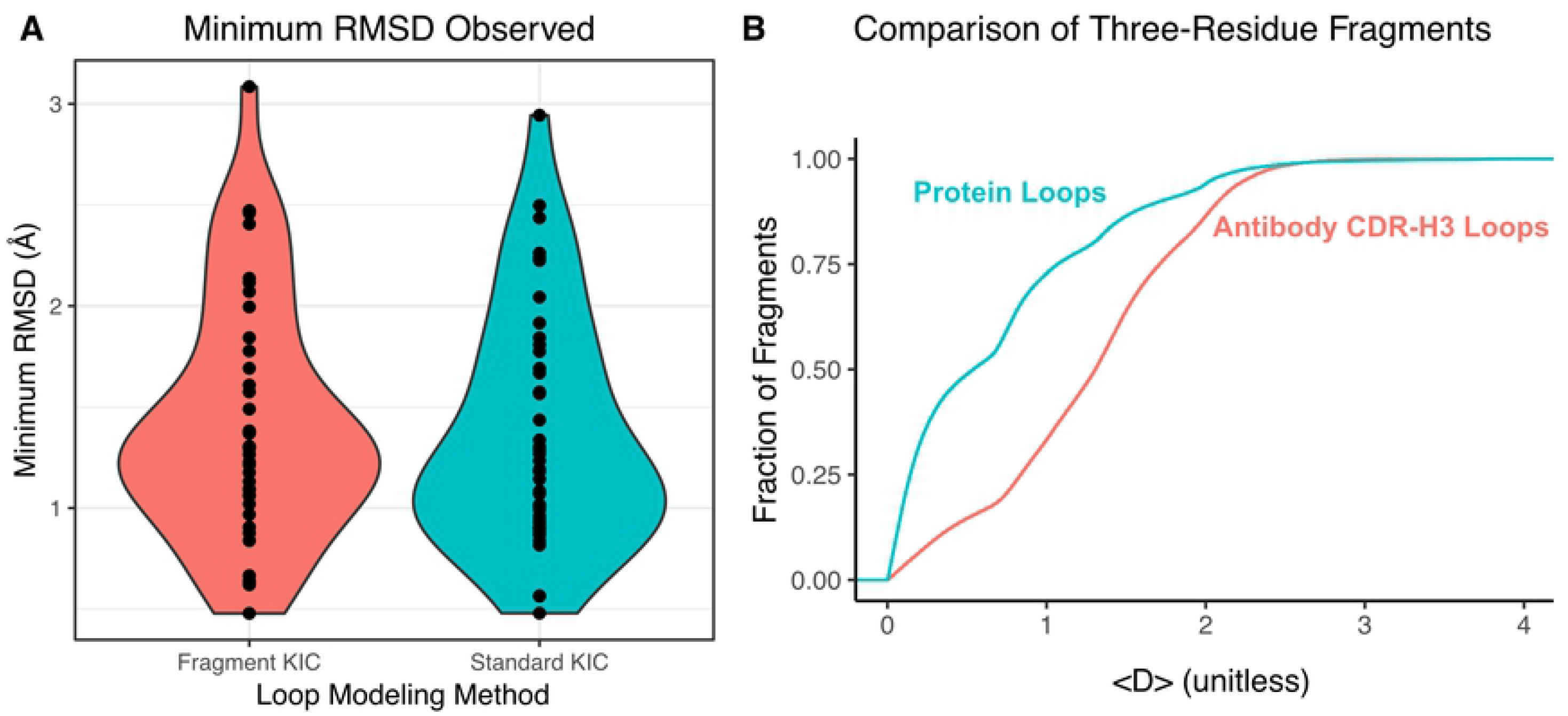
Comparison of loop modeling methods. (A) The distributions of the minimum CDR-H3 loop RMSDs observed for all antibodies in the benchmark, for two loop modeling methods, do not significantly differ according to Student’s t-test (p-value = 0.67). (B) Three-residue fragments from the PDB are more structurally similar to protein loops than to antibody CDR-H3 loops. All three-residue fragments selected by the fragment picker were compared to their corresponding loop sub-regions. For each fragment and loop combination, a chord distance was calculated to compare the difference in dihedral angles: 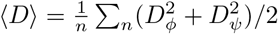 where *D*^2^(*θ*_1_, *θ*_2_) = 2 – 2cos(*θ*_2_ – *θ*_1_). Thus, 〈D〉 has a minimum of 0, if a fragment matches a loop exactly, and a maximum of 4, if a fragment differs by 180 degrees at every dihedral angle. The cumulative distribution function of these distances then yields the probability (y-axis) that a fragment is within a certain chord distance (x-axis) of a loop.

To ensure continuous testing of the CDR-H3 loop modeling stage, we implemented a subset of the Marze *et al.* antibody targets as a scientific benchmark. Specifically, we selected six targets of varying difficulty, based on prior modeling performance [30] and CDR-H3 loop length (Table S1). The scientific benchmark then consists of running the CDR-H3 loop modeling stage on homology models of these antibody frameworks (Appendix S2).

Finally, beyond enabling a new loop modeling approach, we introduced an automated V_H_–V_L_ Q–Q hydrogen bond constraint. Constraints modify the Rosetta score function by adding customizable functions to the standard collection of physical and statistical terms. A typical use case for constraints is to incorporate experimental data in simulations by penalizing protein conformations that are nonconcordant. RosettaAntibody recommends constraining the C-terminal CDR-H3 loop kink and a Q–Q hydrogen bond at the V_H_–V_L_ interface, if present. The kink and the Q–Q hydrogen bond are both present in 81.1% and 88.5% of antibodies in our database. Thus both constraints should be enabled by default. However, the kink constraint was only recently automated [30] and the Q–Q constraint remained user specified until this publication. As a consequence, the constraints were under utilized because they relied on manual user input to identify the corresponding residues and determine the functional form and weights of the constraint.

We implemented a constraint automation similar to the one used by Weitzner and Gray to constrain the kink [30]. Key residues are automatically identified by relying on known sequence features and implementing a consistent numbering scheme throughout modeling. The functional form and weights of the constraint are based on observed geometries in the protein data bank. Using the recently established scientific benchmarking framework, we tested multiple constraint functions and strengths to identify a reasonable default. We found that the harmonic constraint improved the fraction of models in which the hydrogen bond is formed (Figure S3), but did not significantly affect the CDR-H3 loop RMSDs (Figure S4). The constraint is now enabled whenever the requisite glutamine residues are present in the antibody sequence.

### Rosetta SnugDock improvements

The primary improvement to SnugDock was the introduction of a more general `FoldTree` that enabled the modeling of heavy-chain only antibodies. Additionally, we introduced the possibility for fragment-based loop modeling, the capacity for experimental constraints, as well as two automated constraints (as in RosettaAntibody), and scientific benchmarks.

#### FoldTree simplification

Primarily, we improved the kinematics of Rosetta SnugDock. The kinematic layer of Rosetta controls how atomic coordinates are updated over the course of a simulation. It is necessary because Rosetta uses internal coordinates (dihedral angles, with fixed bond lengths and angles) to accelerate sampling in most protocols (simulations in Cartesian coordinates are possible, but not common) [31]. Central to the process of keeping internal and Cartesian coordinates up-to-date is an object known as the `FoldTree`, at the residue level, and the `AtomTree`, at the atomic level [32]. The `FoldTree` is implemented as a directed acyclic graph that propagates coordinates changes. For example, a typical `FoldTree` for a four-protein complex would be linearly ordered, taking the chain order from the PDB file (Fig 1A). In this `FoldTree`, one cannot dock a middle protein independently of its neighbors. This poses a problem in the case of an antibody-antigen complex, where the relative V_H_–V_L_ orientation might change as the antibody accommodates the antigen. This problem is further amplified when modeling loops, as loops require alterations to the `FoldTree` to permit the repeated breaking and closing of covalent bonds. The typical solution is to switch between multiple, incompatible, “simple” `FoldTree` objects that rely on assumptions about the input and have to be specified beforehand. To overcome this issue, we generalized the set of assumptions applied in the `FoldTree` construction stage of SnugDock, resulting in a single, consistent FoldTree that can be used throughout the simulation. This tree also enabled the modeling of heavy-chain only antibodies (*e.g.* camelid).

In the initial implementation of Rosetta SnugDock, it was assumed that the docking partners consisted of a light chain, a heavy chain, and an antigen, in that order. The `FoldTree` was updated at each stage of the simulation to accommodate appropriate sampling. The light chain could be docked to the heavy chain to refine the orientation. In the stage sampling the Ab Ag interface, the `FoldTree` was re-ordered to have the antigen first then the light and heavy chains, so the antigen could be docked to the antibody. Additionally, during H3 and H2 loop modeling stages, a third `FoldTree` was applied to permit opening and closing the loops. This scheme assumed the presence of a light chain, excluding heavy-chain only antibodies from SnugDock simulations.

To correct this issue, we introduced a more hierarchical FoldTree that exploits “virtual” residues – residues that are chemically and physically ignored, but tracked by the `FoldTree` to store positional information. The virtual residues are placed at individual protein and complex centers-of-mass and then connected to corresponding polypeptide chains in a hierarchical fashion (Fig 1B), such that complexes of interest are grouped together (e.g. the two antibody chains or any number of antigen chains). Using virtual residues overcomes the aforementioned challenges. First, by placing the proteins downstream of virtual residues, each chain can have its own internal `FoldTree` without affecting any downstream partner. This permits `FoldTree`-dependent modifications within in each chain (such as loop modeling) to take place, without necessitating a new `FoldTree`. Second, by placing virtual residues at the centers-of-mass of each protein and the relevant complexes, simultaneous docking between multiple partners is now possible in one `FoldTree`. Finally, this `FoldTree` makes no assumptions about the identity of individual chains, so it is compatible with heavy-chain only antibodies.

The new `FoldTree` enabled our participation for Targets 123, 124, and 160 in the blind protein docking challenge called CAPRI, detailed in [23]. Briefly, we ran standard ensemble SnugDock simulations (Appendix S4). The results showed that we were technically able to model the camelid antibodies, but the models were inaccurate due to the challenges associated with modeling longer CDR-H3 loops (11–21 residues).

#### Introducing constraints to SnugDock

We also implemented automatic Q–Q and kink constraints in SnugDock, and further enabled user-defined constraints. Experimental or computationally-derived epitope data (e.g. [33]) can now guide docking. As a proof of principle, we combined hydrogen exchange-mass spectrometry (HX-MS) data with SnugDock. HX-MS measures the backbone amide hydrogen/deuterium exchange rate, and interacting residues, such as those at epitope or paratope, will yield slower exchange rates that can then suggest binding sites for docking. During the docking process, constraints based on pre-processed HX-MS data are applied to the antibody antigen complex. Interactions that satisfy the experimental constraints are rewarded, whereas the interaction that violate the constraints are penalized. We derived a constraint form for each antigen-residue suggested by HX-MS to the closest antibody CDR residue by using the so-called `KofNConstraint` with a flat harmonic potential. A `KofNConstraint` adds the *K* lowest values of a total of *N* constraints to the score, where the *N* constraints are for each residue in the paratope.

As a proof-of-principle, we selected a camelid antibody–ricin complex, 5BOZ [34], to evaluate the utility of constraints. This PDB structure is one of several antibody–ricin complexes for which HX-MS data is available (Weiss, D. D., personal communication). We introduced the data in SnugDock as `KofNConstraints` (Appendix S8). We then ran a local ensemble docking simulation in SnugDock (Appendix S9) and global rigid-body docking simulation with RosettaDock (Appendix S10) [35], both constrained based on the HX-MS data. We found that, when starting from the bound crystal structure, the global search with constraints produced low-scoring (favorable) models of high quality (according to CAPRI criteria), (Figure 5). When using SnugDock and starting from a modeled antibody and unbound antigen crystal structure, constraints did not result in high quality models. Interestingly, these models were able to produce native-like CDR-H3 loop structures. A full study on the utility of constraining antibody–antigen docking simulations with HX-MS constraints is currently in preparation (Zhou, J., Weis, D. D. & Gray, J. J.).

**Fig 5.**
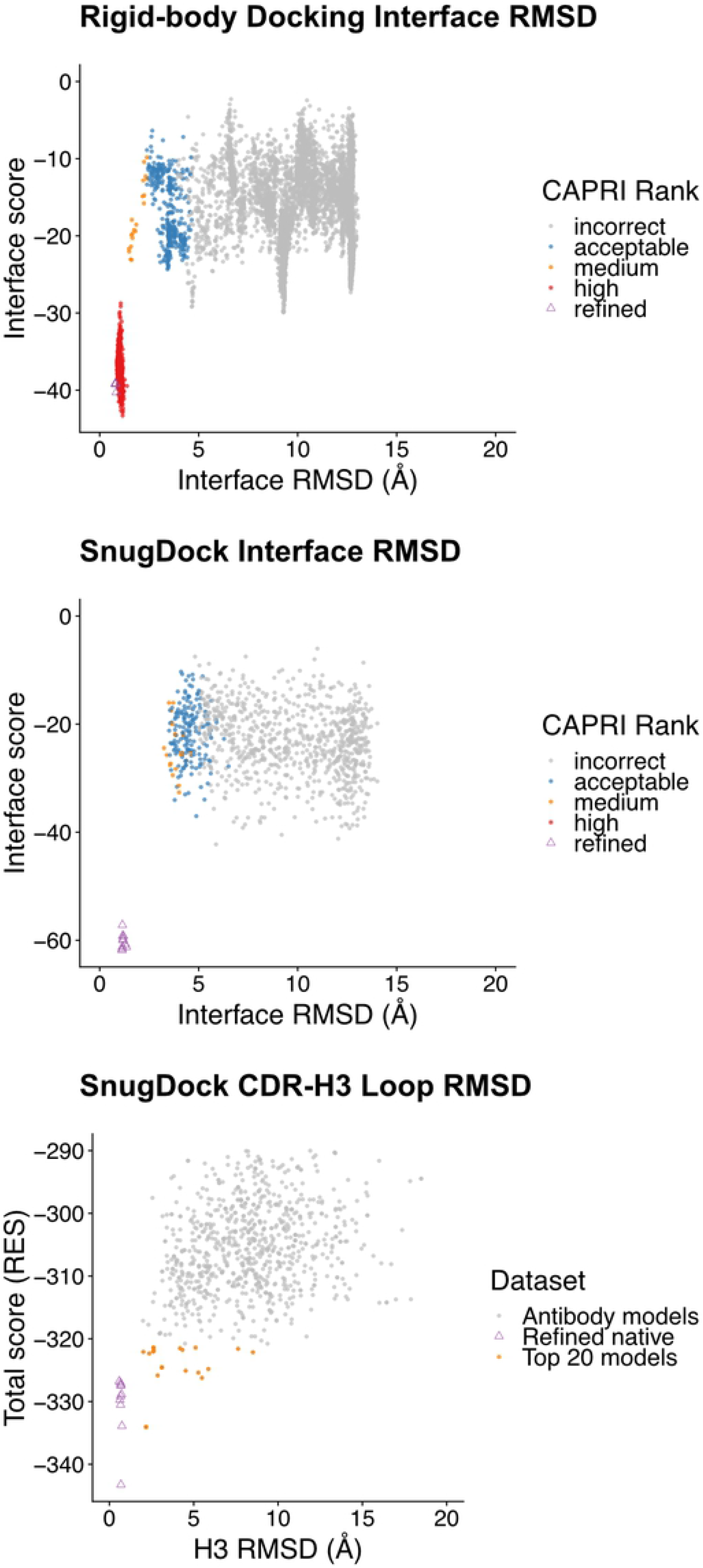
Constraints aid global antibody–antigen docking, but do not affect local refinement. A global, rigid-body search with constraints added to the score function resulted in multiple high-quality models, according to the CAPRI criteria (top panel). However, applying the same constraints to a local search with an antibody homology model did not improve sampling (middle panel). Interestingly, the addition of constraints to SnugDock led to the sampling of native-like CDR-H3 loops, despite not including any constraints during the H3 modeling stage of Rosetta Antibody (bottom panel). “Refined” indicates models created from the bound complex structure, for reference.

## Discussion

Here we presented several advancements in RosettaAntibody and SnugDock that improve performance and collectively lay a foundation for further work. We improved the homology modeling stage of RosettaAntibody by (1) automating the template database to increase coverage and reduce errors, and (2) introducing new filters. We advanced the CDR-H3 loop modeling stage by introducing a new loop modeling approach and structural constraints. We updated SnugDock to use a universal `FoldTree` that enabled the docking of single-domain antibodies, added a new loop modeling, and introduced new constraints. Finally, we implemented scientific benchmarks that regularly test the performance of these protocols.

However, major challenges remain that could be the focus of future development: CDR-H3 modeling, a truly universal `FoldTree` for multi-body docking, and improved selection of non-H3 CDR loop templates. Of these, CDR-H3 loop modeling is the most challenging. Broadly, modeling challenges are binned into two categories: scoring and sampling. We recently showed that native-like antibody loops, when sampled, can be identified by score alone in Rosetta [30]. We also observed that for some targets it is challenging to observe a native-like conformation in the set of all models [6, 23]. Thus, the CDR-H3 loop modeling problem is primarily a sampling challenge. The anecdotal evidence is further supported by observations that CDR-H3 loops are exceptionally diverse, as has been previously demonstrated by others [29] and shown by us here (Figure 4B). One possible approach to overcoming the sampling challenging is to accelerate the loop modeling step to sample more loop conformations. As the slow stage of generating loop models is scoring and filtering, using a knowledge-based rather than physical potential may provide a viable alternative. For example, KORP is a potential capable of scoring 100,000 12-residue loop decoys in under a minute [36]. Another approach to improving sampling would be a more specialized fragment insertion routine during loop modeling. The method used here relied only on sequence similarity to select fragments of either length 3 or 9 and inserted the fragments randomly throughout the loop modeling simulation. An alternative fragment selection approach would not restrict fragment size and might choose fragments from CDR-H3 or H3-like loops. Fragment insertion would focus on the termini that are more structurally conserved regions, *e.g.* approximately 90% of antibodies have a C-terminal “kink”. Finally, emerging deep-learning-based approaches may accelerate CDR-H3 loop sampling. [37] The new loop modeling framework has laid the foundation for exploring further strategies.

The hierarchical `FoldTree` introduced here allows more flexibility in SnugDock and enable the docking of single-domain antibodies. However, true multi-body docking is still not possible as the SnugDock approach is a specialized class, separate from the general docking approach in Rosetta. Moving forward, docking approaches in Rosetta should be unified. *I.e.* the `DockingProtocol` class should be able to provide all docking functionality, based on user specifications and input.

Finally, the homology modeling stage of RosettaAntibody relies on BLAST to select structural templates for query sequences for the various structural regions of an antibody (Table 1). However, most structural regions are small while BLAST is not optimized for aligning short sequences. Thus going forward we must consider alternative approaches to alignment such as custom PSSMs or machine-learning-based approaches [10, 11].

## Conclusion

The role of computational modeling will grow as the throughput of experimental techniques continues to increase. To enable the continued development of the RosettaAntibody and SnugDock protocols, we have simplified their usage, robustified their performance on varied targets, and developed scientific benchmarks. By simplifying the usage of these protocols, future developers can focus on improving the underlying algorithms rather than fiddling with extraneous options. Increasing the utility of these protocols will ensure their longevity as increasingly diverse and challenging pathogens lead to the development and discovery of atypical antibodies. Finally, the availability and regular assessment of scientific benchmarks will encourage a more rapid developmental cycle.

## Supporting information

**Appendix S1.**
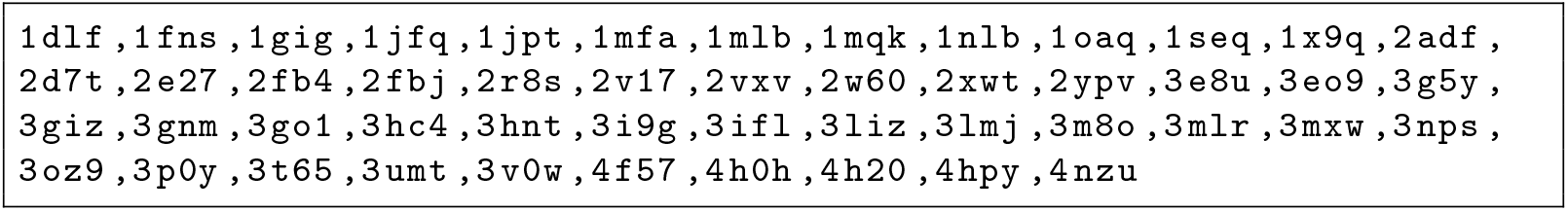
List of antibodies used in the grafting benchmark.

**Appendix S2.**
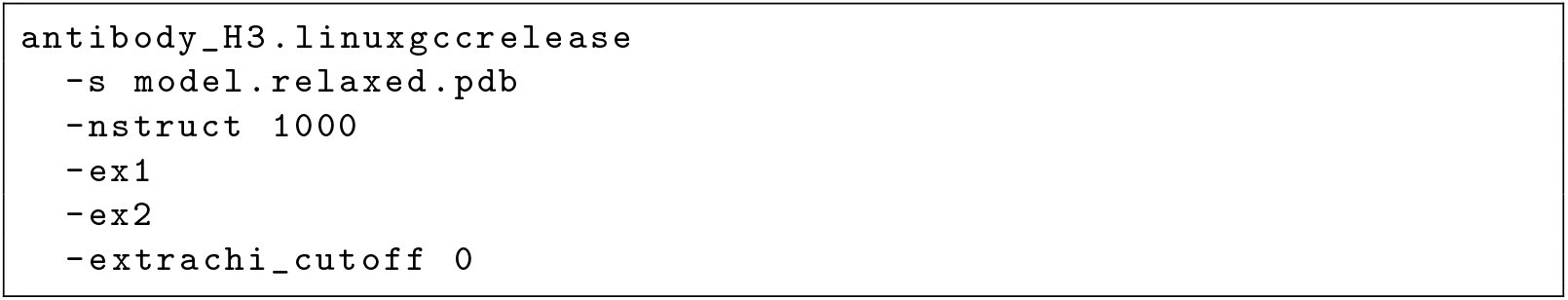
RosettaAntibody command line. Note constraints are now automatically enabled, to disable constraints, use `-antibody:constrain_vlvh_qq false`, `-antibody:h3_loop_csts_lr false` and `-antibody:h3_loop_csts_hr false`.

**Appendix S3.**
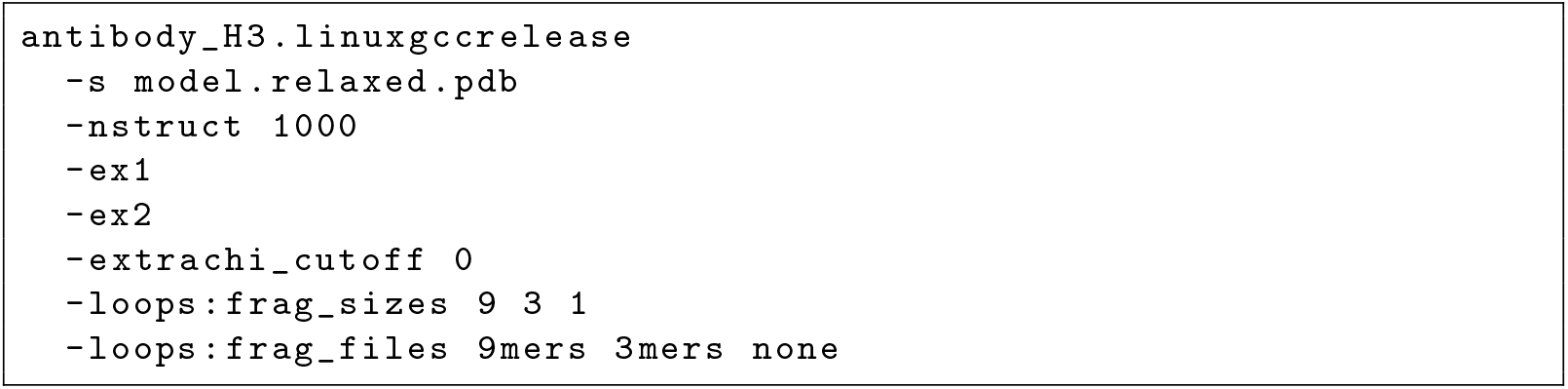
RosettaAntibody command line with fragments. Note constraints are now automatically enabled, to disable constraints, use `-antibody:constrain_vlvh_qq false`, `-antibody:h3_loop_csts_lr false` and `-antibody:h3_loop_csts_hr false`.

**Appendix S4.**
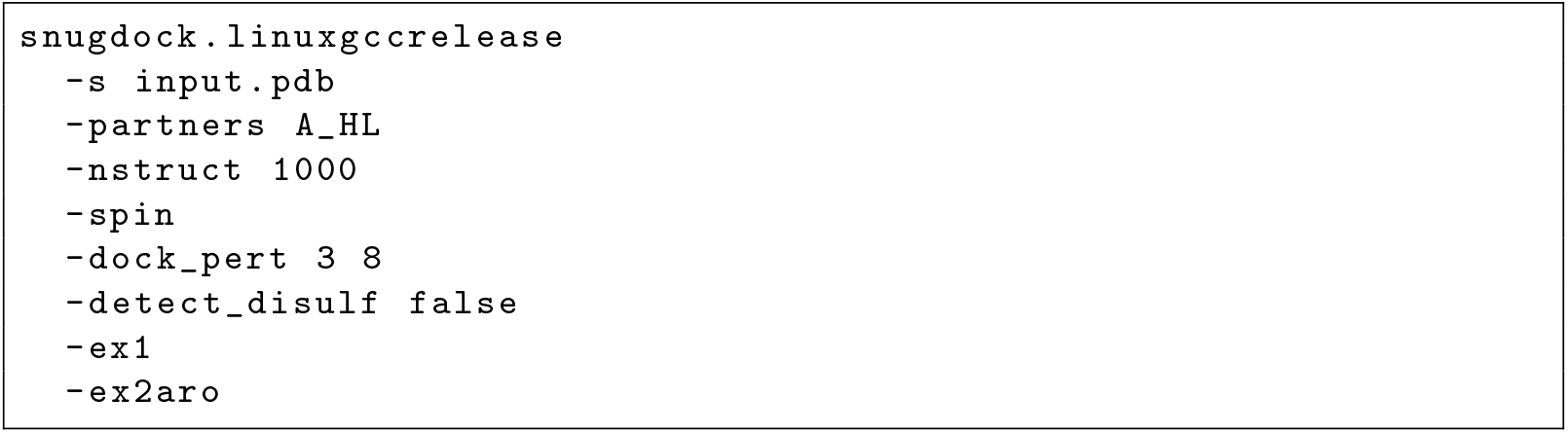
SnugDock command line. Note constraints are now automatically enabled, to disable constraints, use `-antibody:constrain_vlvh_qq false`, `-antibody:h3_loop_csts_lr false` and `-antibody:h3_loop_csts_hr false`.

**Appendix S5.**
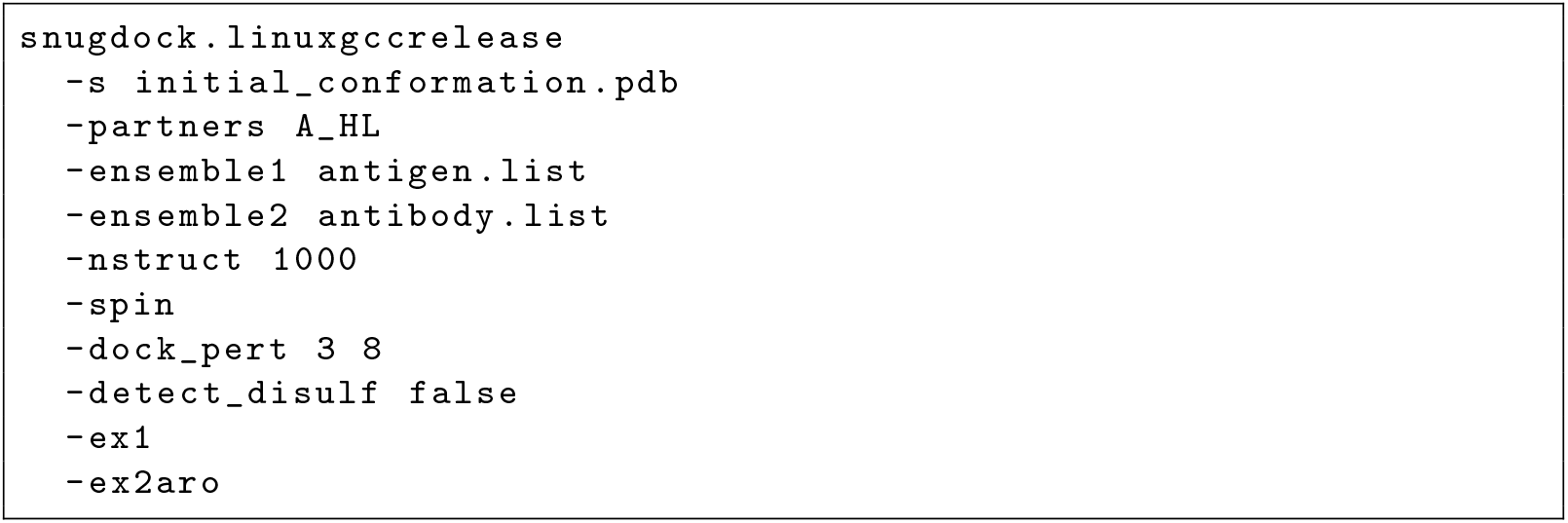
SnugDock command line with an ensemble of structures. Note constraints are now automatically enabled, to disable constraints, use `-antibody:constrain_vlvh_qq false`, `-antibody:h3_loop_csts_lr false` and `-antibody:h3_loop_csts_hr false`. Furthermore, structures must be prepared for ensemble docking by `docking_prepack_protocol` see (below).

**Appendix S6.**
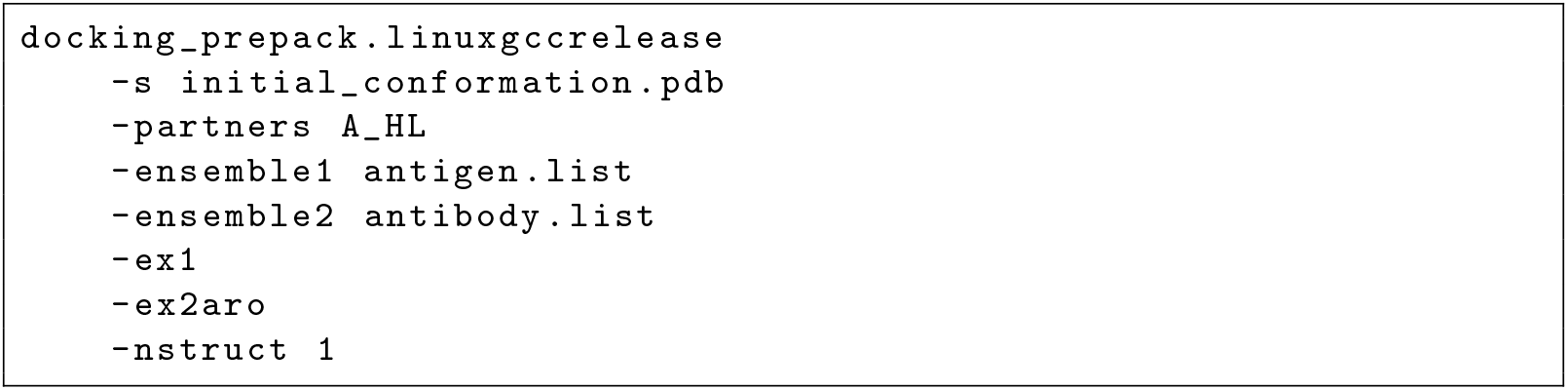
Prepack protocol command line. This will alter the antigen.list and antibody.list files in place. Please note that the chain order in the `-partners` flag must match the order of chains in the PDB passed by the `-s` flag and `-ensemble1` and `-ensemble2`. That is to say in the example below the initial_conformation.pdb file has the “A” chain first followed by “H” and “L” while the first ensemble is a list of antigen only structures and the second ensemble is a list of antibody only structures. All structures must have matching numbers of residues.

**Appendix S7.**
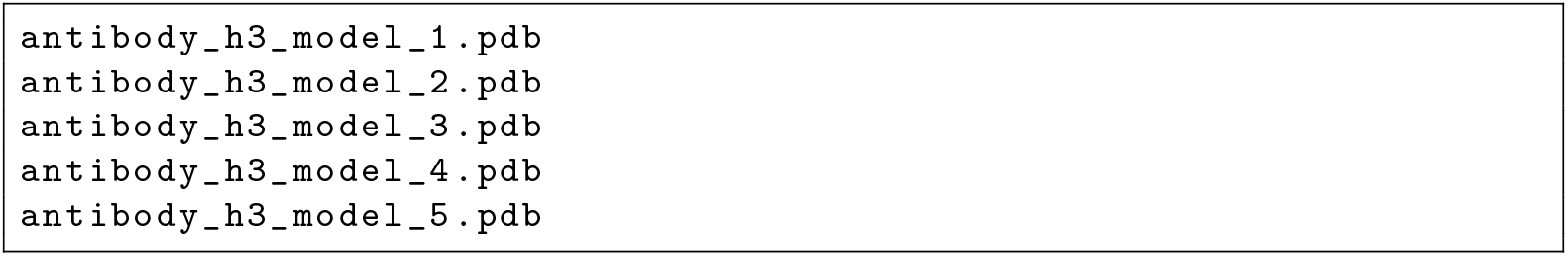
Sample list file. The ensemble of antibody sturctures in this case comes from differ H3 models, but ensembles can also be generated by FastRelax, for example.

**Appendix S8.**
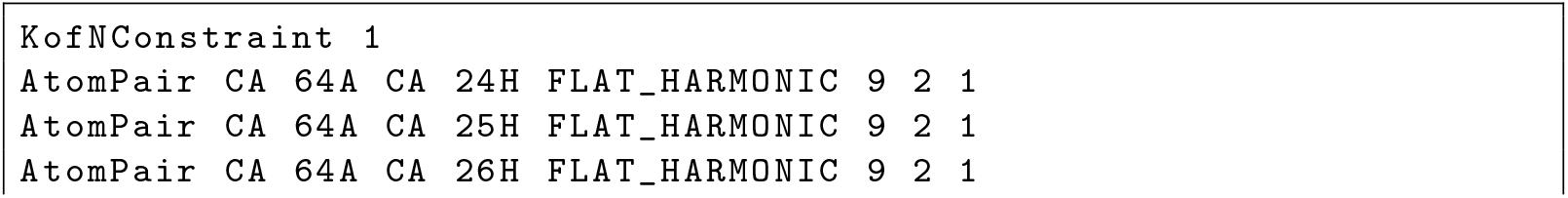
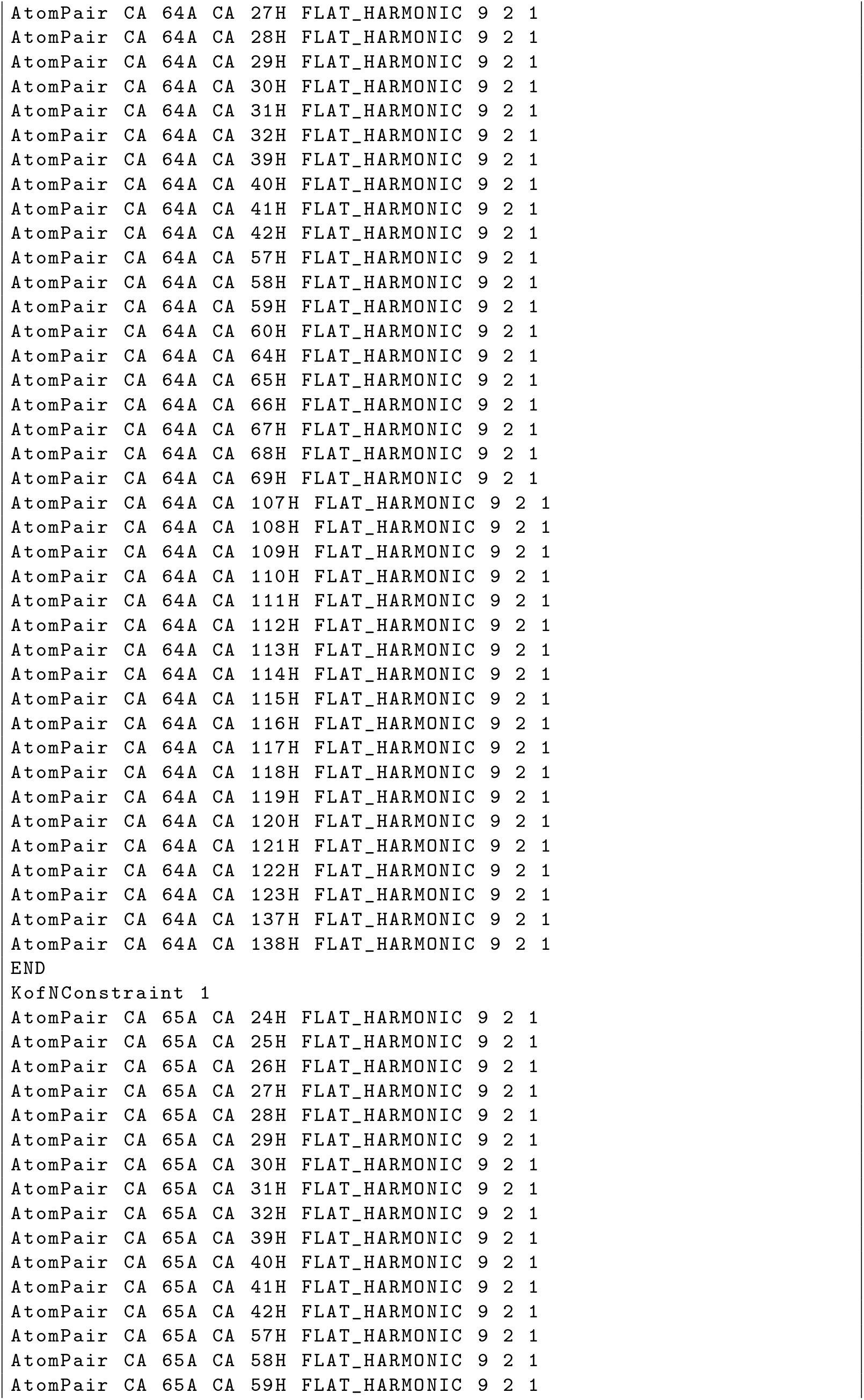
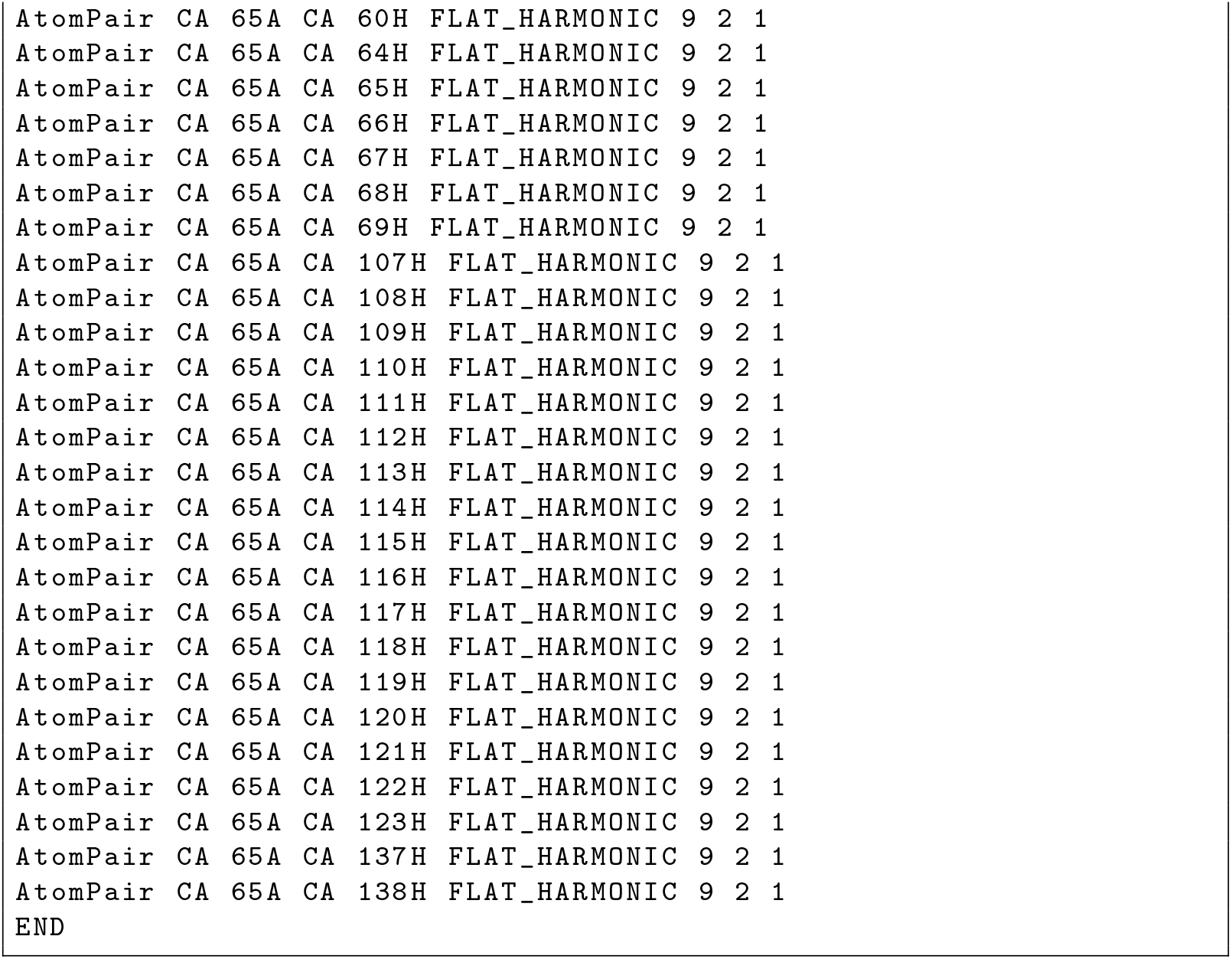
Sample KofNConstraint file. This file only contains two constraints as an example. A complete file would contain one `KofNConstraint` for each antigen residue with HX-MS data. Each `KofNConstraint` would contain one flat harmonic constraint for each CDR residue.

**Appendix S9.**
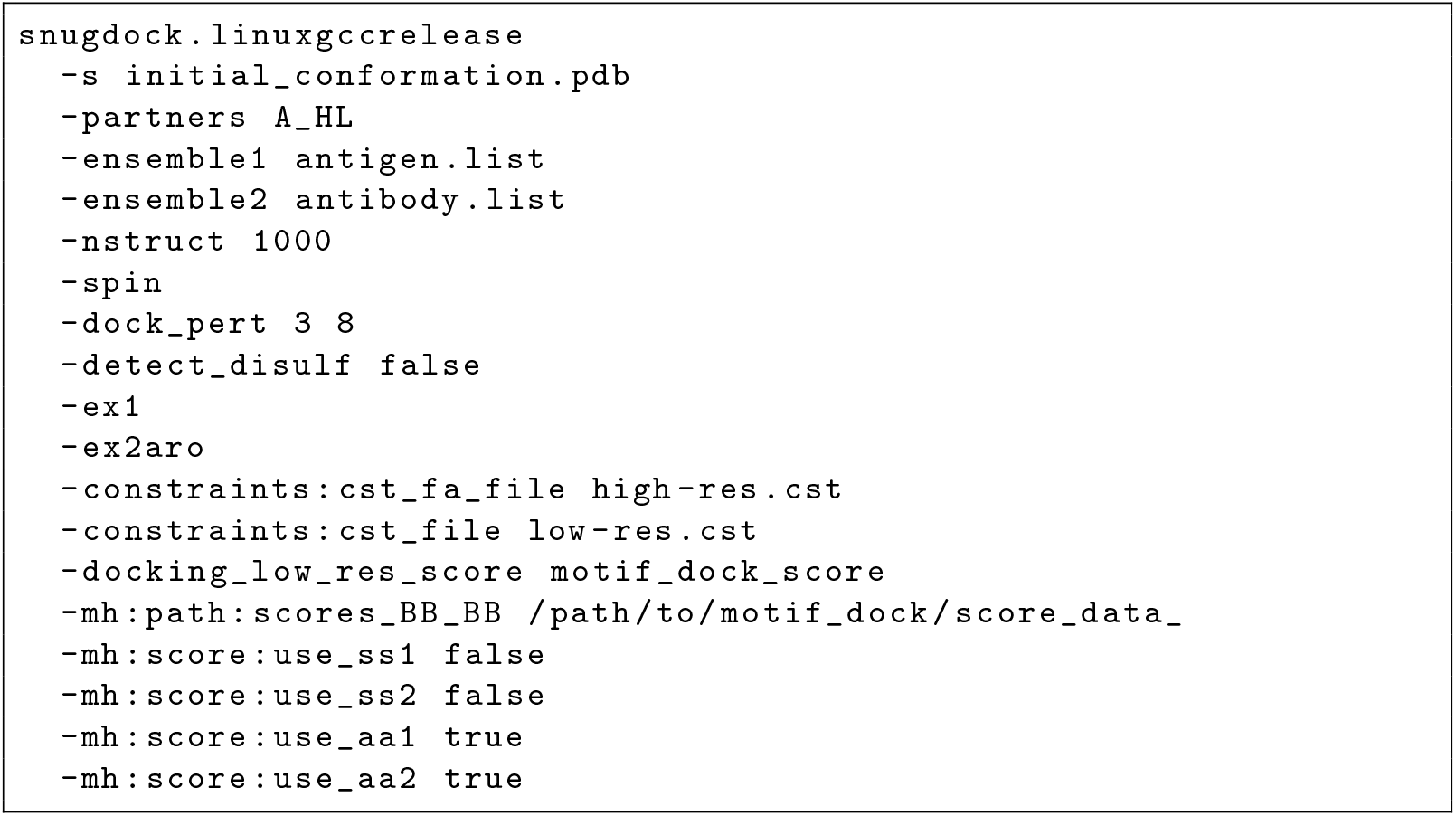
SnugDock command line with constraints and motif dock score (MDS). Additional constraints can be added to both the low- and high-resolution stages of SnugDock. MDS is a special score function for the low-resolution stage of docking. It has been found to improve performance in protein protein complex docking. It can be used in SnugDock as well.

**Appendix S10.**
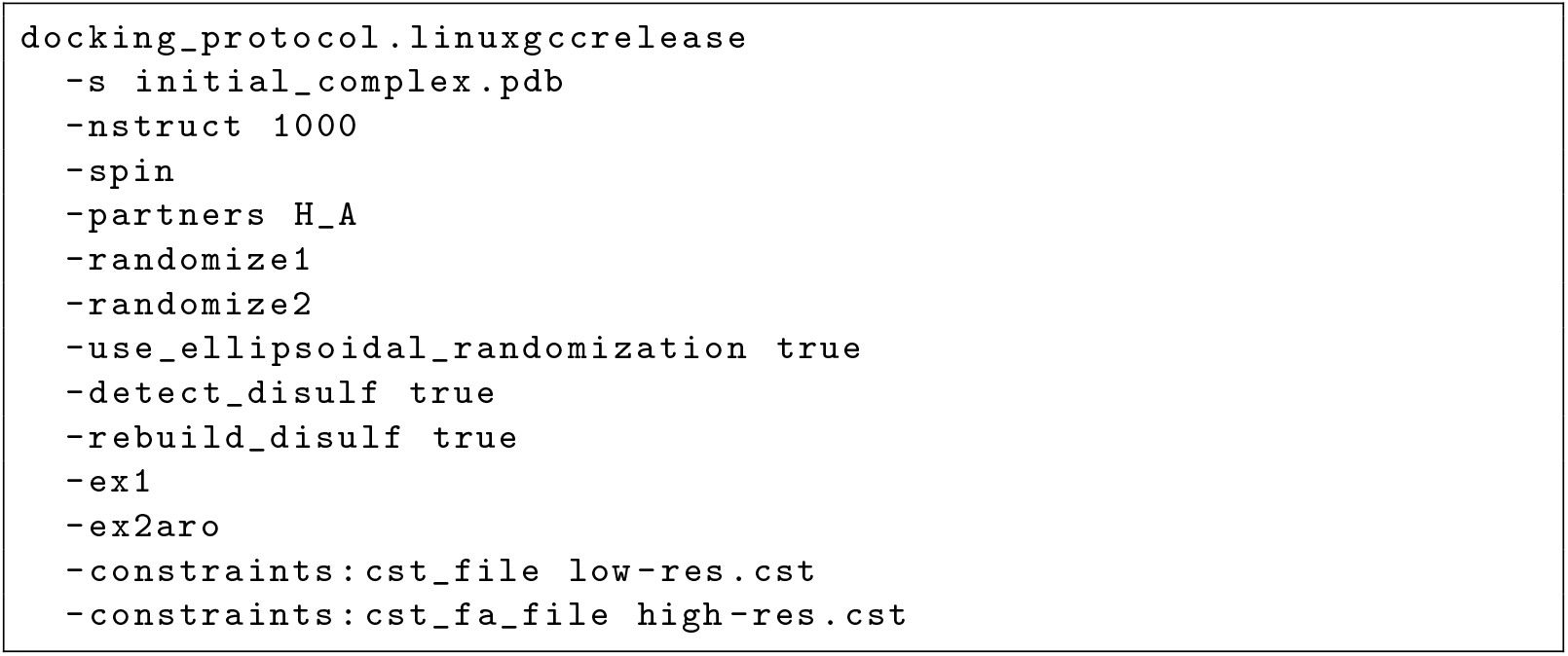
Global docking command line. Exemplary flags for global docking with constraints.

**Table S1.** Target antibody CDR-H3 loops for the antibody modeling scientific benchmark.

| PDB ID | CDR-H3 Length | Difficulty |
| --- | --- | --- |
| 1DLF | 12 | Easy |
| 4HPY | 13 | Easy |
| 2VXV | 14 | Medium |
| 3M8O | 10 | Medium |
| 1SEQ | 16 | Hard |
| 4HPY | 18 | Hard |

**Table S2.**
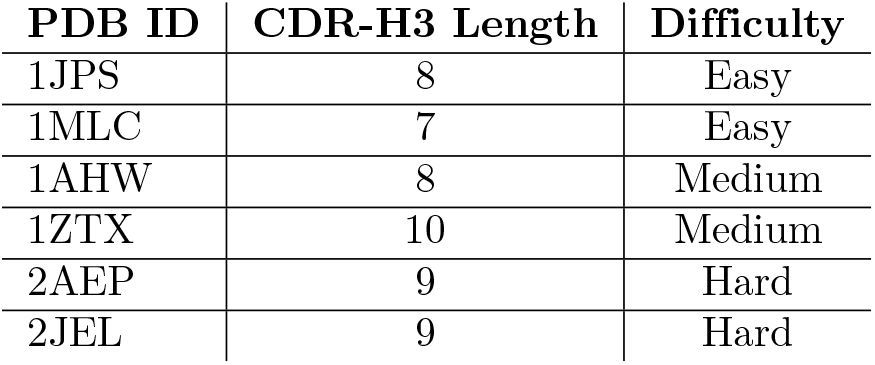
Target antibody–antigen complexes for the docking scientific benchmark.

**Figure S1.**
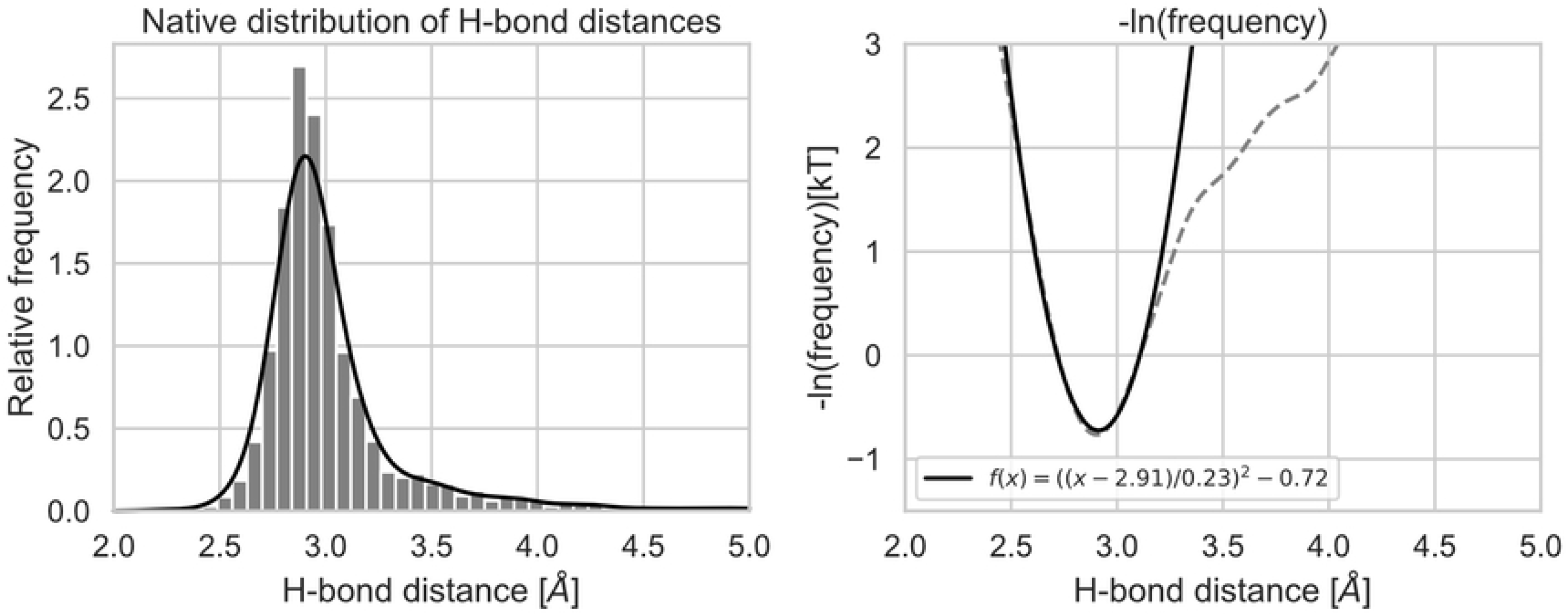
Q–Q hydrogen bond distances observed in the RosettaAntibody database. **Left:** The histogram depicts the observed distances between the oxygen and nitrogen atoms of light chain residue Q38 and heavy chain residue Q39. The distribution was fit by kernel density estimate using Gaussian kernels. **Right:** The negative logarithm of the probability is proportional to the energy. A harmonic function was fit in the range of 2.5 A to 3.1 Å.

**Figure S2.**
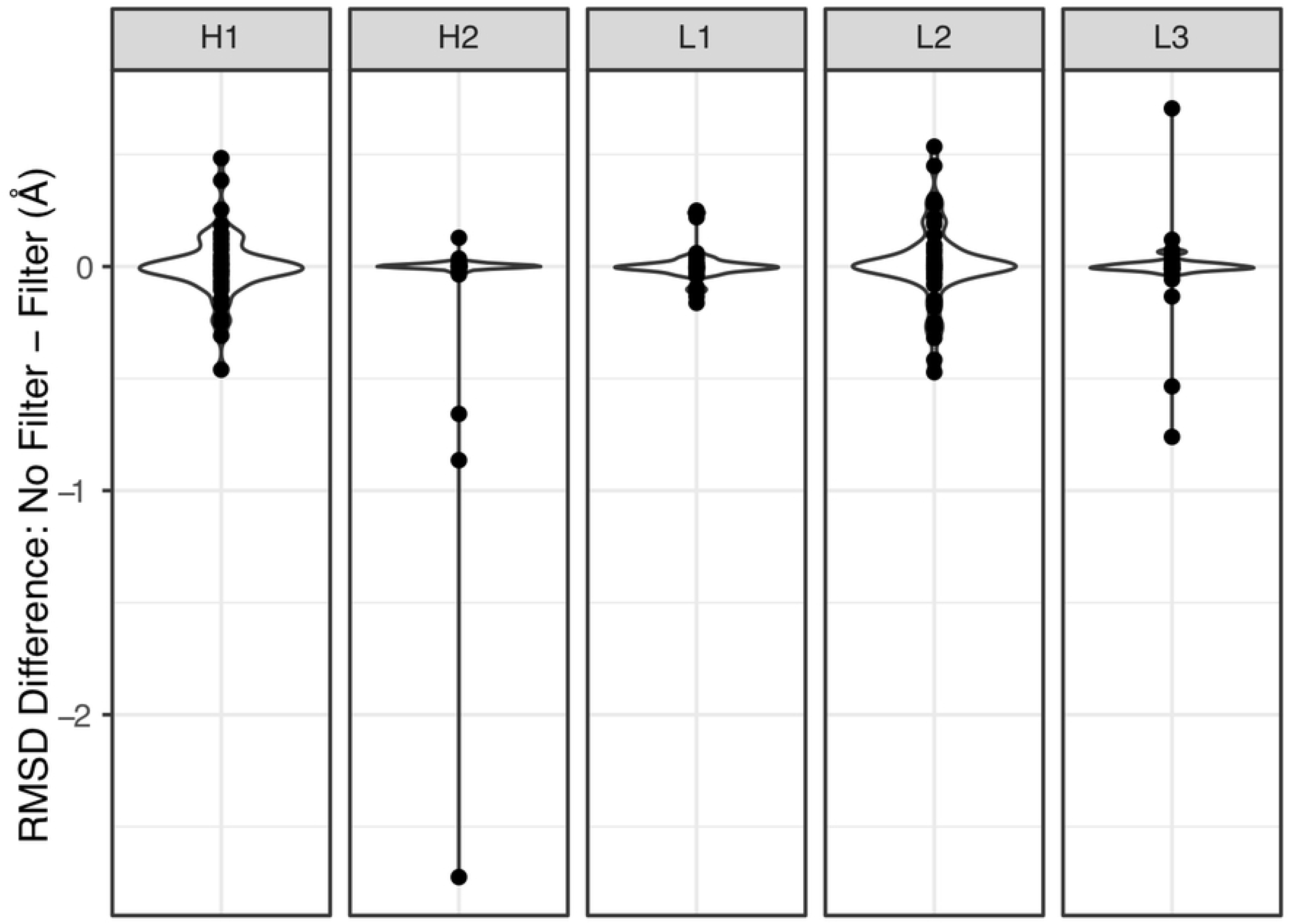
Proline filter has minimal effect on grafted model RMSDs. Comparison of the non-H3 CDR loop RMSDs before and after the application of a proline filter. The filter prevents the use of a template when there is a mismatched proline residue with the query. The differences show that most loops are unaffected. In one case for the CDR H2 loop, the loop is model is worse following the application of the filer (moving 2 Å further from the native). This is exclusively due to the presence of an glycine at the start of the target loop (PDB ID: 3LMJ). In the initial model (PDB ID 6EIK, no proline filter), the template also has a glycine, correctly modeling the initial loop structure, whereas the proline-filter-selected template (PDB ID 5LSP) lacks this initial glycine and cannot accurately model the loop start resulting in a cascading worsening of the loop model. All other loops show minor variations within 1 Å.

**Figure S3.**
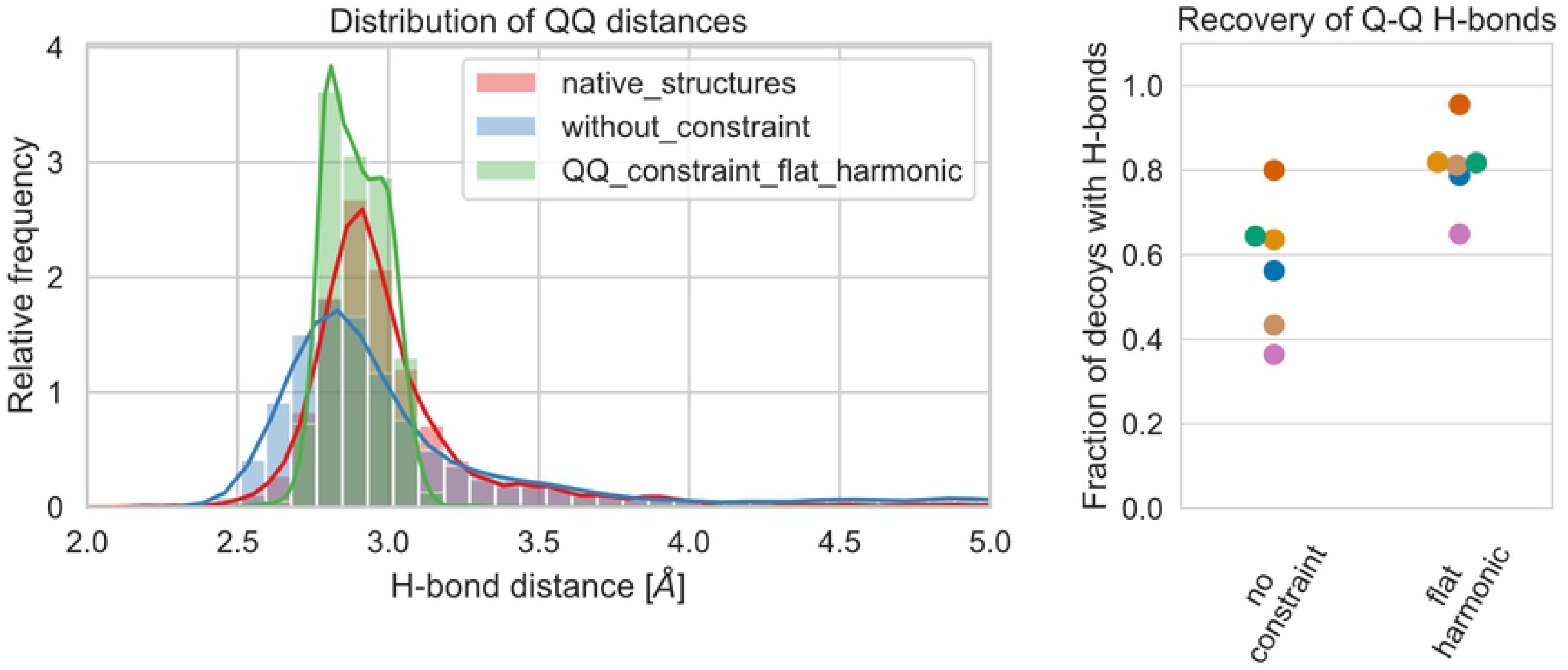
The Q–Q constraint increases the fraction of models that form hydrogen bonds. We generated 500 decoys of 6 antibodies with solved structures (Table S1) either without or with a flat harmonic constraint between the relevant Gln residues. Left: The distances between the nitrogen and oxygen atoms of residues Q38 of the light chain and Q39 of the heavy chain were measured and compared to the native distributions in our antibody database. Right: Each decoy was analyzed for presence of the two possible hydrogen bonds using PyRosetta’s get_hbonds() function. The fraction of decoys forming both hydrogen bonds is shown for each antibody (color-coded).

**Figure S4.**
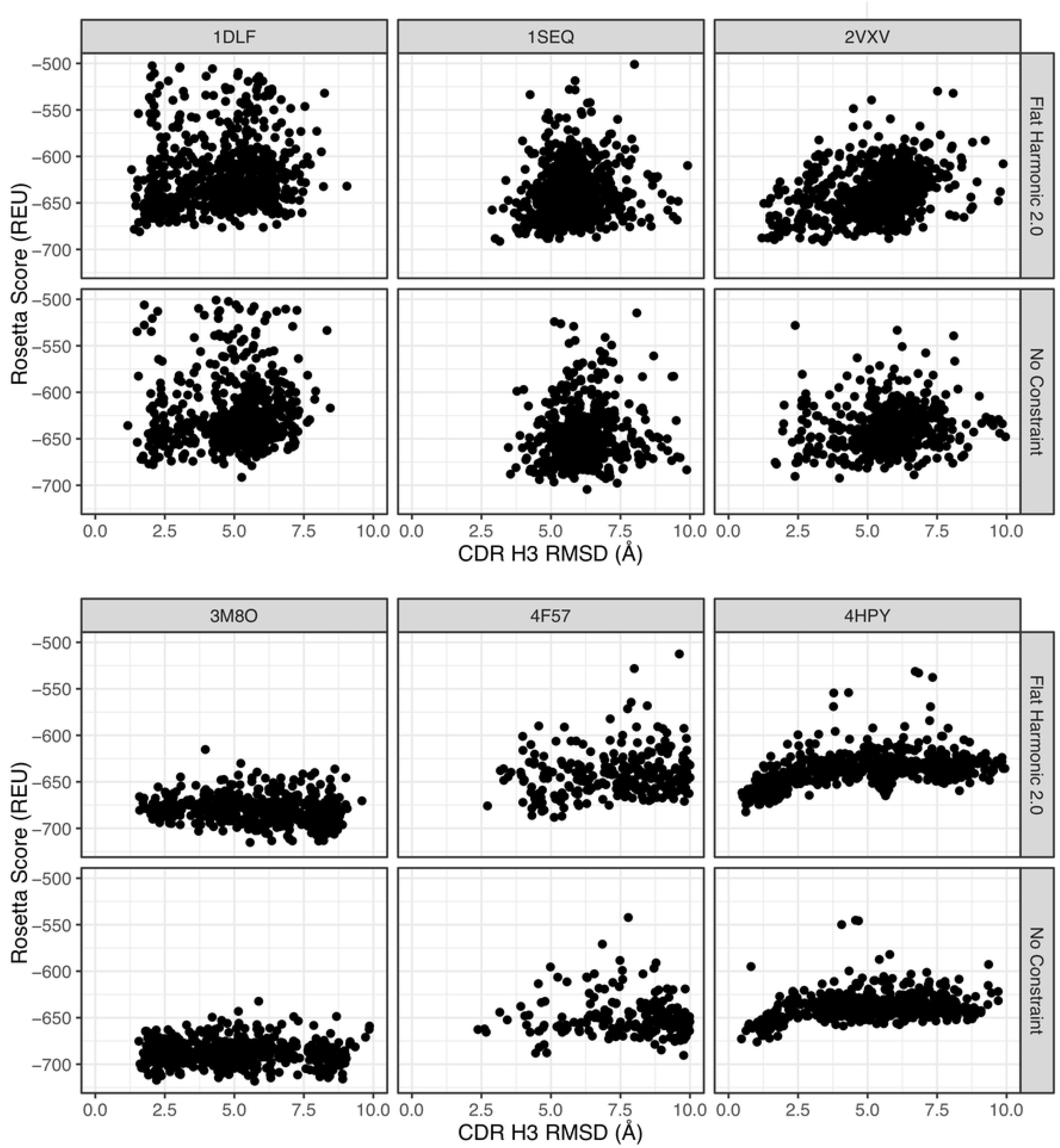
The Q–Q constraint does not appear to have a strong effect on CDR-H3 loop modeling. A funnel plot (total score versus CDR-H3 loop RMSD) comparison of RosettaAntibody on six benchmark antibodies does not show a significant difference after the incorporation of a flat harmonic. The constraint seemingly improves performance on targets 2VXV and 4F57, but worsens it on 3M8O.

## Acknowledgments

The authors of this study would like to acknowledge Drs. Nicholas Marze and Brian Weitzner (both of Johns Hopkins University) for helpful discussions and advice, Xingjie Pan and Prof. Tanja Kortemme (both of University of California San Francisco) for help with implementing the refactored loop modeling code for antibody modeling, and Dr. Julia Koehler Leman (Flatiron Institute) and Sergey Lyskov (Johns Hopkins University) for implementing the benchmarking server. Prof. David Weis (University of Kansas) advised the work including HX-MS data.

## References

1. Georgiou G, Ippolito GC, Beausang J, Busse CE, Wardemann H. Quake SR. The promise and challenge of high-throughput sequencing of the antibody repertoire. Nat. Biotechnol. 2012;32: 158–168.

2. Krawczyk K, Kelm S, Kovaltsuk A, Galson JD, Kelly D, Trueck J, et al. Structurally mapping antibody repertoires. Front. Immunol. 2018;9: 1698

3. Leem J, Dunbar J, Georges G, Shi J, Deane CM. ABodyBuilder: automated antibody structure prediction with data–driven accuracy estimation. mAbs 2016;8: 1259–1268.

4. Lepore R, Olimpieri PP, Messih MA, Tramontano A. PIGSPro: prediction of immunoglobulin structures v2. Nucleic Acids Research 2017;45: W17–W23.

5. Weitzner BD, Jeliazkov JR, Lyskov S, Marze N, Kuroda D, Frick R, et al. Modeling and docking of antibody structures with Rosetta. Nat. Protoc. 2016;12: 401–416.

6. Weitzner BD, Kuroda D, Marze N, Xu J, Gray JJ. Blind prediction performance of RosettaAntibody 3.0: Grafting, relaxation, kinematic loop modeling, and full CDR optimization. Proteins 2014;82: 1611–1623.

7. Almagro JC, Teplyakov A, Luo J, Sweet RW, Kodangattil S, Hernandez-Guzman F, et al. Second antibody modeling assessment (AMA-II). Proteins 2014;82: 1553–1562.

8. North B, Lehmann A, Dunbrack Jr. RL. A new clustering of antibody CDR loop conformations. J. Mol. Biol. 2011;406: 228–256.

9. Al-Lazikani B, Lesk AM, Chothia C. Standard conformations for the canonical structures of immunoglobulins. J. Mol. Biol. 1997;273: 927–948.

10. Long X, Jeliazkov JR, Gray JJ. Non-H3 CDR template selection in antibody modeling through machine learning. PeerJ 2019;7: e6179; doi: 10.7717/peerj.6179.

11. Wong WK, Georges G, Ros F, Kelm S, Lewis AP, Taddese B, Leem J, et al. SCALOP: sequence-based antibody canonical loop structure annotation. Bioinformatics 2019;35: 1774–1776.

12. James LC, Roversi P, Tawfik DS. Antibody multispecificity mediated by conformational diversity. Science 2003;299: 1362–1367.

13. Kozakov D, Hall DR, Xia B, Porter KA, Padhorny D, Yueh C, et al. The ClusPro web server for protein-protein docking. Nat. Protoc. 2017;12: 255–278.

14. Brenke R, Hall DR, Chuang GY, Comeau SR, Bohnuud T, Beglov D, et al. Application of asymmetric statistical potentials to antibody–protein docking. Bioinformatics 2012;28: 2680–2614.

15. Ramírez-Aportela E, López-Blanco JR, Chacón P. FRODOCK 2.0: fast protein-protein docking server. Bioinformatics 2016;32: 2386–2388.

16. Mashiach E, Schneidman-Duhovny D, Peri A, Shavit Y, Nussinov R, Wolfson HJ. An integrated suite of fast docking algorithms. Proteins 2010;78: 3197–3204.

17. Sircar A, Gray JJ. SnugDock: paratope structural optimization during antibody–antigen docking compensates for errors in antibody homology models. PLoS Comput. Biol. 2010;6: e1000644; doi: 10.1371/journal.pcbi.1000644.

18. Méndez R, Leplae R, De Maria L, Wodak SJ. Assessment of blind predictions of protein-protein interactions: Current status of docking methods. Proteins 2003;52: 51–67.

19. Guest JD, Vreven T, Zhou J, Moal I, Jeliazkov JR, Gray JJ, et al. An expanded benchmark for antibody–antigen docking and affinity prediction reveals insights into antibody recognition determinants. 2020; Preprint at SSRN: https://ssrn.com/abstract=3564997.

20. Dunbar J, Krawczyk K, Leem J, Baker T, Fuchs A, Georges G, et al. SAbDab: the structural antibody database. Nucleic Acids Research 2014;42: D1140–D1146.

21. Chothia C, Lesk AM. Canonical structures for the hypervariable regions of immunoglobulins. J Mol Biol. 1987;196: 901–917.

22. Marze NA, Lyskov S, Gray JJ. Improved prediction of antibody VL-–VH orientation. Protein Engineering, Design and Selection 2016;29: 409–418.

23. Roy Burman SS, Nance ML, Jeliazkov JR, Labonte JW, Lubin JH, Biswas N, et al. Novel sampling strategies and a coarse-grained score function for docking homomers, flexible heteromers, and oligosaccharides using Rosetta in CAPRI Rounds 37–45. Proteins 2019; https://doi.org/10.1002/prot.25855.

24. Kaas Q, Ruiz M, Lefranc MP. IMGT/3Dstructure-DB and IMGT/StructuralQuery, a database and a tool for immunoglobulin, T cell receptor and MHC structural data. Nucleic Acids Research 2004;32: D208–D210.

25. Swindells MB, Porter CT, Couch M, Hurst J, Abhinandan KR, Nielsen JH, et al. abYsis: integrated antibody sequence and structure—management, analysis, and prediction. J. Mol. Bio. 2017;429: 356–364.

26. Abhinandan KR, Martin ACR. Analysis and improvements to Kabat and structurally correct numbering of antibody variable domains. Molecular Immunology 2008;45: 3832–3839.

27. Stein A, Kortemme T. Improvements to robotics-inspired conformational sampling in Rosetta. PLOS ONE 2013;8: e63090; doi: 10.1371/journal.pone.0063090.

28. Gront D, Kulp DW, Vernon RM, Strauss CEM, Baker D. Generalized fragment picking in Rosetta: design, protocols and applications. PLOS ONE 2011;6: e23294; doi: 10.1371/journal.pone.0023294.

29. Regep C, Georges G, Shi J, Popovic B, Deane CM. The H3 loop of antibodies shows unique structural characteristics. Proteins 2017;85: 1311–1318 (2017).

30. Weitzner BD, Gray JJ. Accurate structure prediction of CDR H3 loops enabled by a novel structure-based C-terminal constraint. J. Immunol. 2016; doi: 10.4049/jimmunol.1601137.

31. Parsons J, Holmes JB, Rojas JM, Tsai J, Strauss CE. Practical conversion from torsion space to Cartesian space for in silico protein synthesis. J. Comput. Chem. 2005;26: 1063–1068.

32. Wang C, Bradley P, Baker D. Protein–Protein docking with backbone flexibility. JMB 2007;373: 503–519.

33. Hua, C. K., et al. Computationally-driven identification of antibody epitopes. eLife 2017;6: e29023; doi: 10.7554/eLife.29023.

34. Rudolph MJ, Vance DJ, Cassidy MS, Rong Y, Mantis NJ. Structural analysis of single domain antibodies bound to a second nneutralizing hot spot on ricin toxin’s enzymatic subunit. J. Biol. Chem. 2017;292: 872–883.

35. Marze NA, Roy Burman SS, Sheffler W, Gray JJ. Efficient flexible backbone protein–protein docking for challenging targets. Bioinformatics 2018;34: 3461–3469.

36. López-Blanco JR, Chacón P. KORP: knowledge-based 6D potential for fast protein and loop modeling, Bioinformatics 2019;35: 3013–3019.

37. Ruffolo JA, Guerra C, Mahajan SP, Sulam J, Gray JJ. Geometric Potentials from Deep Learning Improve Prediction of CDR H3 Loop Structures. bioRxiv https://doi.org/10.1101/2020.02.09.940254 (2020).

